# Elevated METTL9 is associated with peritoneal dissemination in human scirrhous gastric cancers

**DOI:** 10.1101/2021.09.13.460036

**Authors:** Toshifumi Hara, Yuuki Tominaga, Koji Ueda, Keichiro Mihara, Kazuyoshi Yanagihara, Yoshifumi Takei

**Affiliations:** Department of Medicinal Biochemistry, School of Pharmacy, Aichi Gakuin University, 1-100 Kusumoto-cho, Chikusa-ku, Nagoya 464-8650, Japan; Project for Personalized Cancer Medicine, Cancer Precision Medicine Center, Japanese Foundation for Cancer Research, 3-8-31 Ariake, Koto-ku, Tokyo 135-8550, Japan; Department of International Center for Cell and Gene Therapy, Fujita Health University, 1-98 Dengakugakubo, Kutsukake-cho,Toyoake 470-1192, Japan; Division of Translational Research, Exploratory Oncology and Clinical Trial Center, National Cancer Center Hospital East, 6-5-1 Kashiwanoha, Kashiwa 277-8577 Japan

**Keywords:** scirrhous gastric cancer, peritoneal dissemination, methyltransferase-like (METTL) gene, short hairpin RNA, molecular targeting therapy against cancer metastasis

## Abstract

Methylation, the most common chemical modification of cellular components such as DNA, RNA, and proteins, impacts biological processes including transcription, RNA processing, and protein dynamics. Although abnormal expression of methyltransferase can lead to various diseases including cancers, little is known about the relationship between methyltransferase and cancers. Here we aimed to understand the role of methyltransferase in cancer metastasis. We found that elevated methyltransferase-like 9 (METTL9) is closely associated with the acquisition of metastatic activity in human scirrhous gastric cancers. The stable knockdown of METTL9 via an shRNA vector technique in our original metastatic cells from scirrhous gastric cancer patients significantly inhibited migration and invasion. In metastatic cells, METTL9 protein is predominantly localized in mitochondria, and the METTL9 knockdown significantly reduced mitochondrial Complex I activity. METTL9 can be a promising molecular target to inhibit peritoneal dissemination of scirrhous gastric cancers. This report is the first to describe the relationship between METTL9 and cancer metastasis.

**Highlights:**

- Elevated METTL9 correlates with metastasis in human scirrhous gastric cancer.
- This is the first report on the biological relationship between METTL9 and metastasis.
- METTL9 protein localizes mainly in mitochondria in metastatic scirrhous gastric cancer.
- METTL9 knockdown reduces mitochondrial Complex I activity to decrease cell migration and invasion in metastatic scirrhous gastric cancer.
- METTL9 holds promise against peritoneal dissemination of scirrhous gastric cancer.

## 1. Introduction

Adaptation of cells to various environmental and physiological conditions is essential for cell survival and development. In a phenomenon called cell reprogramming, cells quickly change their transcriptomic and metabolic dynamics to adapt to extracellular conditions [1, 2]. Cell reprogramming is achieved in part by chemical modifications such as methylation, acetylation, and phosphorylation [3, 4]. Chemical modifications of DNA, RNA, and proteins substantially affect the stability, activity, and functions of cell components [5]. Furthermore, the aberrant regulation of chemical modification leads to the onset of diverse diseases, including cancers [6].

Human scirrhous gastric cancers are a poorly differentiated and aggressive type of gastric cancer that frequently show peritoneal dissemination, leading to poor prognosis [7, 8]. For patients harboring peritoneal dissemination, effective therapeutics have not yet been established because the molecular mechanisms of peritoneal dissemination of the cancer are not fully understood. To elucidate the molecular mechanisms underlying peritoneal dissemination, we previously established patient-derived metastatic cell lines in human scirrhous gastric cancers [9]. By analyzing metastatic cell models, we discovered the expression alteration of methyltransferase-like (METTL) genes. The METTL gene family consists of 27 members, some of which are known as representative writers of N^6^-methyladenosine (m^6^A), which is the most abundant chemical modification in an RNA molecule [10]. It is known that dysregulation of METTL3 induces abnormal m6A abundance, which plays important roles in the progression of various cancers [11]. However, the functions of most METTL family genes, including cancer metastasis, remain unclear. So far, the biological functions of METTL9 have been exhibited only in autosomal recessive deafness 22 by genetic analysis [12]. Here we aim to investigate the functions of METTL9 genes in the peritoneal dissemination (peritoneal metastasis) of human scirrhous gastric cancer using our metastatic cell model and clinical samples (gastric cancer tissues) from patients. As a result, we discovered that elevated METTL9 expression can function to positively promote peritoneal dissemination in scirrhous gastric cancers.

## 2. Material and methods

### 2.1. Cells

HSC-58 cells (parental cells) were originally established from a patient with scirrhous gastric cancer [9]. 58As9 cells (metastatic cells) were successfully isolated by repeating orthotopic implantation of HSC-58 in nude mice, as described previously [9]. These cell lines were maintained in PRMI-1640 medium (Sigma-Aldrich Japan, Tokyo, Japan) containing 10% fetal bovine serum (FBS) at 37°C in a 5% CO_2_ incubator.

### 2.2. Short hairpin RNA (shRNA) expression vector

For the METTL9 knockdown, an SGEP vector provided by Dr. Johannes Zuber was used [13]. We used the splashRNA algorithm [14] to select target sequences for METTL9 according to the instructions. As a negative control, shRNA against a Renilla luciferase was used. After the introduction of the SGEP-based vectors into 58As9 cells (metastatic cells), the cells were cultured in the presence of puromycin at 4 μg/ml for 1 week. Further, the largest portion of GFP-positive cells (<10%) was isolated by an SH800 Cell Sorter (SONY, Tokyo, Japan). The target sequences of the two shRNAs for human METTL9 were as follows:

shMETTL9# 1: 5’-TTAAGTATAAAAAATATCTTCC-3’.
shMETTL9#2: 5’-TTAAATTCCTAC ATGATATTTA-3’.

Information on the sequences of shRNAs targeting the human METTL9 gene is summarized in Supplementary Table 1.

### 2.3. Antibodies

We purchased anti-human METTL9 antibody (15120-1-AP) from Proteintech Japan (Tokyo, Japan) and anti-β-actin antibody (A5441) from Sigma-Aldrich Japan (Tokyo, Japan).

### 2.4. Reverse transcription quantitative PCR (RT-qPCR)

To isolate total RNAs, ISOGEN reagent (Nippon Gene, Tokyo, Japan) was used according to the manufacturer’s instructions. Normal human stomach tissue RNAs were purchased from TaKaRa Bio (Shiga, Japan). iScript RT Supermix for RT-qPCR (Bio-Rad Laboratories, Hercules, CA, USA) was employed for the synthesis of first-strand cDNA. qPCR was carried out with the StepOne Real-time PCR System (ThermoFisher Scientific). The relative expression level was calculated by the ddCq method using the level of the reference gene, succinate dehydrogenase complex subunit A (SDHA). The sequences of the primer sets used in this study were as follows:

SDHA forward, 5’-TGGGAACAAGAGGGCATCTG-3’
SDHA reverse, 5’-CCACCACTGCATCAAATTCATG-3’
METTL9 forward, 5’-CTGTGATCAGCCCCTGACTT-3’
METTL9 reverse, 5’-TGATGGTTTCTCCCACTTGCC-3’

Information on the sequences of specific primers to determine human METTL9 gene expression is summarized in Supplementary Table 2.

### 2.5. Immunoblotting

Total cell lysates were isolated using RIPA buffer (ThermoFisher Scientific) in the presence of a protease inhibitor mixture (Roche) according to the manufacturer’s instructions. Protein concentrations were determined using the Pierce BCA Protein Assay Kit (ThermoFisher Scientific). The cell extracts were separated by SDS-PAGE and blotted onto Immobilon-P membranes (ATTO, Tokyo, Japan). The membranes were blocked with 5% skim milk in Trisbuffered saline with 0.1% Tween 20 (TBS-T) and further incubated with specific antibodies. The membranes were then incubated with horseradish peroxidase-conjugated secondary antibodies (Cell Signaling Technology, Tokyo, Japan). The specific band was detected using the ECL Prime Western Blotting Detection Reagent (GE Healthcare, Chicago, IL, USA) with the ImageQuant LAS 4000 digital imaging system (GE Healthcare).

### 2.6. Cell proliferation assay

Cell proliferation was examined using Cell Counting Kit-8 (CCK-8; Dojindo, Kumamoto, Japan) according to the manufacturer’s instructions. Briefly, cells were seeded in a 96-well plate (2×10^3^ cells/well). After 24, 48, and 72 hours of seeding, CCK-8 solution was added to each well, and absorbance at 450 nm was measured with a Spark plate reader (TECAN Japan, Tokyo, Japan).

### 2.7. Cell migration and invasion assay

Cells were cultured in the absence of FBS for 24 hours prior to the cell invasion assay. For the assay, a trans-well insert (8 μm pore size) was coated with Matrigel (BD Biosciences, Franklin Lakes, NJ, USA) for at least 2 hours. Cells (2 × 10^5^) were plated onto the top chamber in medium in the absence of FBS. The lower chamber was filled with 0.5 ml medium containing 10% FBS as a chemoattractant. After 48 hours of incubation, the cells that did not migrate or invade through the pores were removed with a cotton swab. Cells on the lower surface of the membrane were fixed and stained with crystal violet. Pictures of the membrane were taken by using a digital camera, and the invaded cells were counted.

### 2.8. Immunofluorescence staining

Cells were cultured for 1 day on round coverslips (Fisher Scientific) in a 12-well plate. For staining of mitochondria, MitoTracker^™^ Red CMXRos (ThermoFisher Scientific) was used according to the manufacturer’s instructions. Cells were fixed with 4% paraformaldehyde in PBS and then treated with 0.1% Triton X-100 in PBS for permeabilization. The fixed cells were treated with Blocking One Histo (Nacalai Tesque) and then incubated with anti-human METTL9 antibody (1:200 dilution ratio) for 1 hour at room temperature. Further, an anti-rabbit IgG antibody conjugated with Alexa Fluor 455 (ThermoFisher Scientific) was added. The nuclei were stained with Cellstain DAPI solution (Dojindo). Images were obtained with a confocal laser scanning fluorescence microscope, LSM 800 with Airyscan (Zeiss, Jena, Germany).

### 2.9. Mitochondrial activity assay

Mitochondrial fractions from the cells were isolated by the conventional method. The protein concentration of a fraction was determined by the Pierce BCA Protein Assay Kit. An equal amount of mitochondrial fraction was subjected to the MitoCheck Complex I Activity Assay Kit (Cayman Chemical, Ann Arbor, MI, USA). The Complex I activity was determined by calculating the slope of the initial linear portion of the curve according to the manufacturer’s instructions.

### 2.10. Resources from a public database

For analysis of the prognostic relevance of METTL9, we used a Kaplan-Meier plotter for gastric cancer patients with poor differentiation (n=166).

### 2.11. Clinical samples

The experiments using patients’ samples were performed with Institutional Review Board approval from Aichi Gakuin University and the National Cancer Center Research Institute. We obtained appropriate informed consent from the patients with gastric cancers in accordance with the Declaration of Helsinki. The cancer tissues were harvested, and snap frozen until used for the experiments. The RNA extraction methods were described previously [9].

### 2.12. Statistical analysis

All experiments were performed in duplicate at least three times, and representative data were shown. Statistical analysis was performed by the paired t-test. A p-value less than 0.05 was considered to be statistically significant. The Spearman’s rank correlation value was analyzed by Coloc 2, a FIJI plugin.

## 3. Results

### 3.1. Elevated METTL9 is associated with peritoneal dissemination of scirrhous gastric cancers

METTL9 expression was significantly elevated in our metastatic 58As9 cells compared with HSC-58 cells (parental cells) via RT-qPCR (Fig. 1A). We also showed very low expression of METTL9 in normal stomach tissues from healthy donors (Fig. 1A). Similar to the results for RT-qPCR, the METTL9 protein level was increased in metastatic 58As9 cells (Fig. 1B) compared with HSC-58 cells (parental cells). METTL9 expression was significantly elevated in cancer tissues from patients with scirrhous gastric cancers (Fig. 1C). In particular, in primary gastric cancer tissues from patients with peritoneal dissemination (Fig. 1C, third black bar) METTL9 expression was significantly higher than that in primary gastric cancer tissues from patients without peritoneal dissemination (Fig. 1C, second white bar). METTL9 expression was further significantly increased in the peritoneal disseminated cancer tissues (Fig. 1C, fourth black bar). These results suggest that elevated METTL9 expression accompanies cancer progression of scirrhous gastric cancers and that METTL9 must play some important role(s) in peritoneal dissemination in scirrhous gastric cancers. Finally, using a public database, we analyzed the prognostic relevance of METTL9 gene expression in patients with poorly differentiated gastric cancers (Fig. 1D). According to this analysis, a high expression level of METTL9 is significantly associated with poor prognosis (Fig. 1D).

**Fig. 1.**
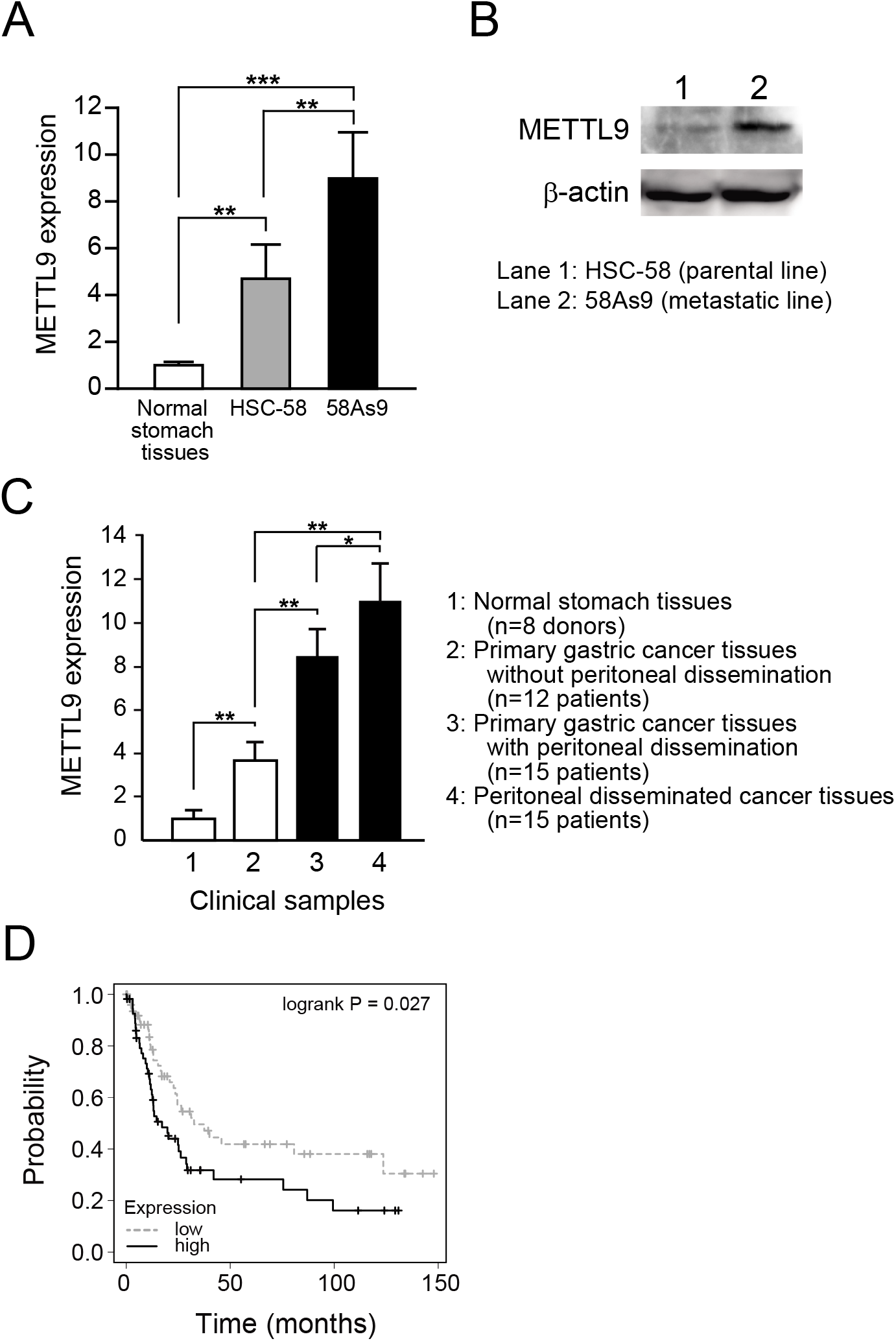
Elevated METTL9 expression is correlated with metastatic activity of human scirrhous gastric cancers. (A) Expression levels of METTL9 by RT-qPCR in parental cells HSC-58, metastatic cells 58As9, and normal human stomach tissues. **, p<0.01; and ***, p<0.001. (B) The protein level of METTL9 was determined by immunoblotting with specific antibodies. (C) Expression levels of METTL9 by RT-qPCR in clinical samples from gastric cancer patients. *, p<0.05; and **, p<0.01. (D) Overall survival of patients with poorly differentiated gastric cancer was analyzed by splitting the levels of METTL9 expression among patients (n=166). The statistical significance of patient survival was observed between patients with low METTL9 expression and those with high METTL9 expression (p=0.027).

The series of impressive results in Fig. 1 prompted us to explore the role of METTL9 in peritoneal dissemination in scirrhous gastric cancers. We were also motivated because the functions of METTL9 in cancer development and progression had not been elucidated yet.

### 3.2. Knockdown of METTL9 expression significantly reduces metastatic activity in scirrhous gastric cancers

To understand the cellular functions of METTL9 particularly in metastatic activity, we successfully generated 58As9 METTL9 knockdown cells by introducing shRNA-expressing vectors (Fig. 2). 58As9 METTL9 knockdown cells showed markedly reduced METTL9 expression at both the mRNA and protein levels (Fig. 2A and 2B) compared with the levels in 58As9 cells introduced with control shRNA (shRluc). The 58As9 METTL9 knockdown cells showed no morphological changes (data not shown). Knockdown of METTL9 expression in 58As9 cells significantly decreased cell proliferation (Fig. 2C), migration via trans-well assay (Fig. 2D), and invasion via Matrigel invasion assay (Fig. 2E) compared with the control 58As9 cells (shRluc). These data suggest that elevated METTL9 expression contributes to the development and progression in scirrhous gastric cancers.

**Fig. 2.**
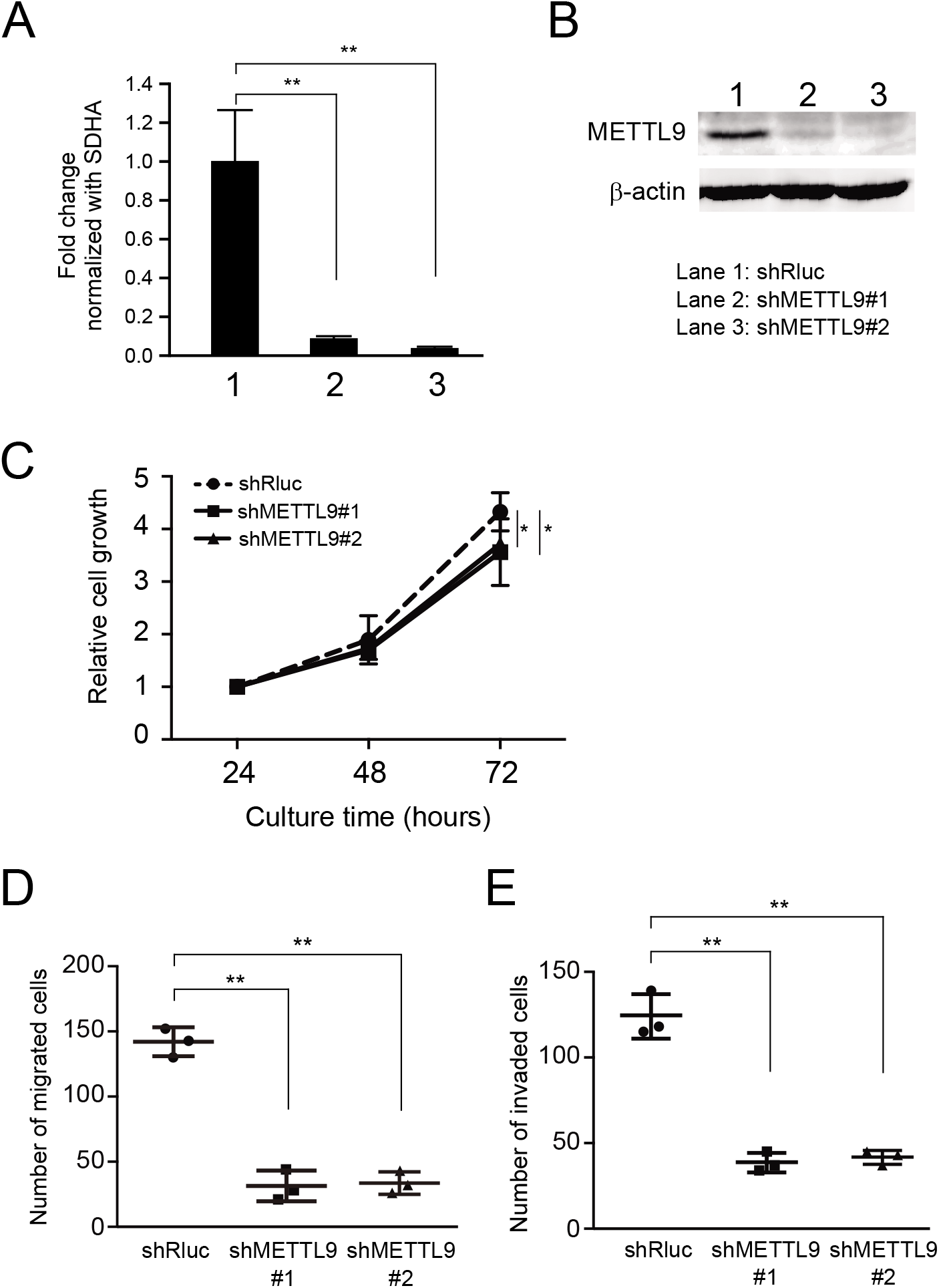
Knockdown of METTL9 reduces cell proliferation, migration, and invasion in 58As9 metastatic cells. (A) The knockdown effect of shRNAs targeting human METTL9 was examined by RT-qPCR. Two shRNAs (shMETTL9#1 and shMETTL9#2) were used independently. METTL9 shRNA-introduced 58As9 cells showed significantly reduced levels of METTL9 expression. **, p<0.01. 1, shRluc (control); 2, shMETLL9#1; and 3, shMETTLE9#2. (B) The knockdown (protein levels of METTL9) was assessed by immunoblotting. (C) The effect of METTL9 knockdown on cell proliferation was analyzed. METTL9 knockdown significantly attenuated cell proliferation. *, p<0.05. (D) METTL9 knockdown in 58As9 significantly reduced cell migration via transwell assay. **, p<0.01. (E) METTL9 knockdown in 58As9 significantly reduced cell invasion activity. **, p<0.01.

In our previous report [9], we established another cell line pair from another scirrhous gastric cancer patient: HSC-44PE parental cells, and 44As3 metastatic cells. The latter were established from the former via the orthotopic transplantation method as well [9]. In 44As3, we observed elevated METTL9 expression via RT-qPCR, and we tried to knockdown METTL9 also in 44As3 cells. Both shMETTL9#1 and shMETTL9#2 showed significantly reduced METTL9 expression compared with shRluc (Supplementary Fig. 1A). Using the METTL9-knockdown 44As3 cells, we successfully showed a significant inhibition of cell proliferation (Supplementary Fig. 1B, clonogenic assay) and migration (Supplementary Fig. 1C, woundhealing assay) compared with the control cells (shRluc). According to these control experiments using another metastatic line, 44As3, we found that elevated METTL9 is generally associated with peritoneal dissemination in scirrhous gastric cancer. We also found that elevated METTL9 in metastatic cells serves to maintain the cells’ motility, which is a driving force behind peritoneal dissemination.

### 3.3. METTL9 protein is predominantly colocalized with the mitochondrial compartment in our metastatic cells

Using 58As9 cells, the subcellular distribution of METTL9 protein was examined via immunofluorescence staining and confocal microscopic analysis. We found that METTL9 protein was predominantly localized in the mitochondria (Fig. 3A). To explore METTL9 protein subcellular distribution in greater detail, we further analyzed the intensity-based spatial correlation. Pearson’s rank correlation value between METTL9 and mitochondria was 0.928. In addition, the intensity profiling of METTL9 well matched that of mitochondria (Fig. 3C). We also found that a small amount of METTL9 was partially localized in the nucleus (Fig. 3B and 3C). These results are consistent with a recent finding that METTL9 mediates 1-methylhistidine modification mainly targeting the component of mitochondrial respiratory Complex I [15]. Taken together, the previous and present results indicate that METTL9 can affect the functions of mitochondrial respiratory Complex I in our metastatic 58As9 cells.

**Fig. 3.**
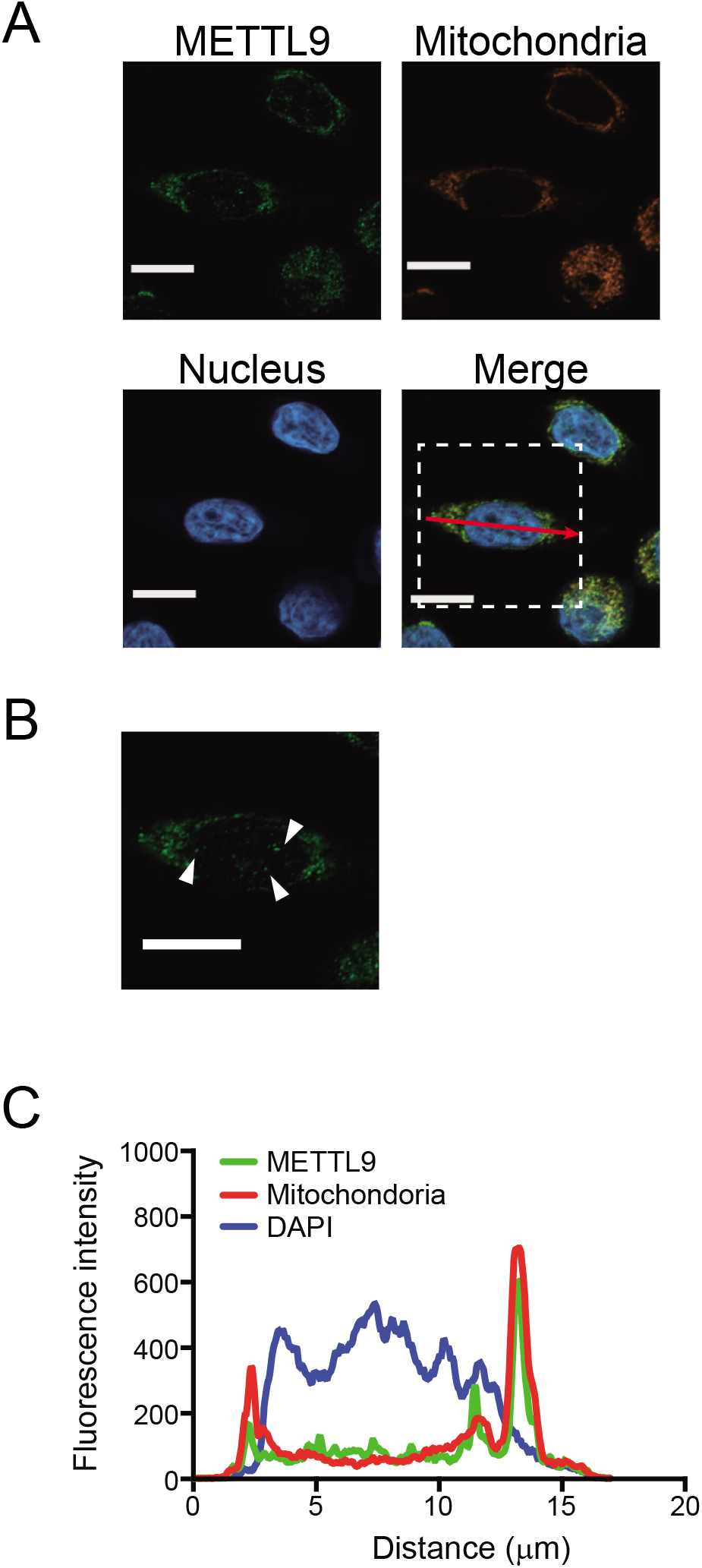
METTL9 protein localizes on mitochondria in 58As9 metastatic cells. (A) Subcellular distribution of METTL9 protein was examined by an immunostaining approach. 58As9 cells were stained with anti-METTL9 antibody and secondary antibody with green fluorescence. Their nuclei were stained with 4’,6-diamidino-2-phenylindole (DAPI). Mitochondria were detected with MitoTracker Red CMXRos reagent. Scale bars, 10 μm. (B) Enlargement of the dashed square from (A). White arrowheads indicate METTL9 localization in the nucleus, which is independent of the mitochondrial region. Scale bar, 10 μm. (C) The intensity profile shown by the red arrow in (A).

### 3.4. METTL9 affects mitochondrial Complex I activity in our metastatic cells

To examine whether or not METTL9 affects mitochondrial Complex I, we measured the rate of NADH oxidation using mitochondrial fractions isolated from METTL9-knockdown 58As9 cells (Fig. 4A). We found a two-fold reduction in mitochondrial Complex I activity in METTL9-knockdown 58As9 cells compared with the control cells (Fig. 4B). These results indicate that METTL9-mediated 1-methylhistidine modifications are important in mitochondrial activity and can affect the acquisition of metastatic activity of cancer cells.

**Fig. 4.**
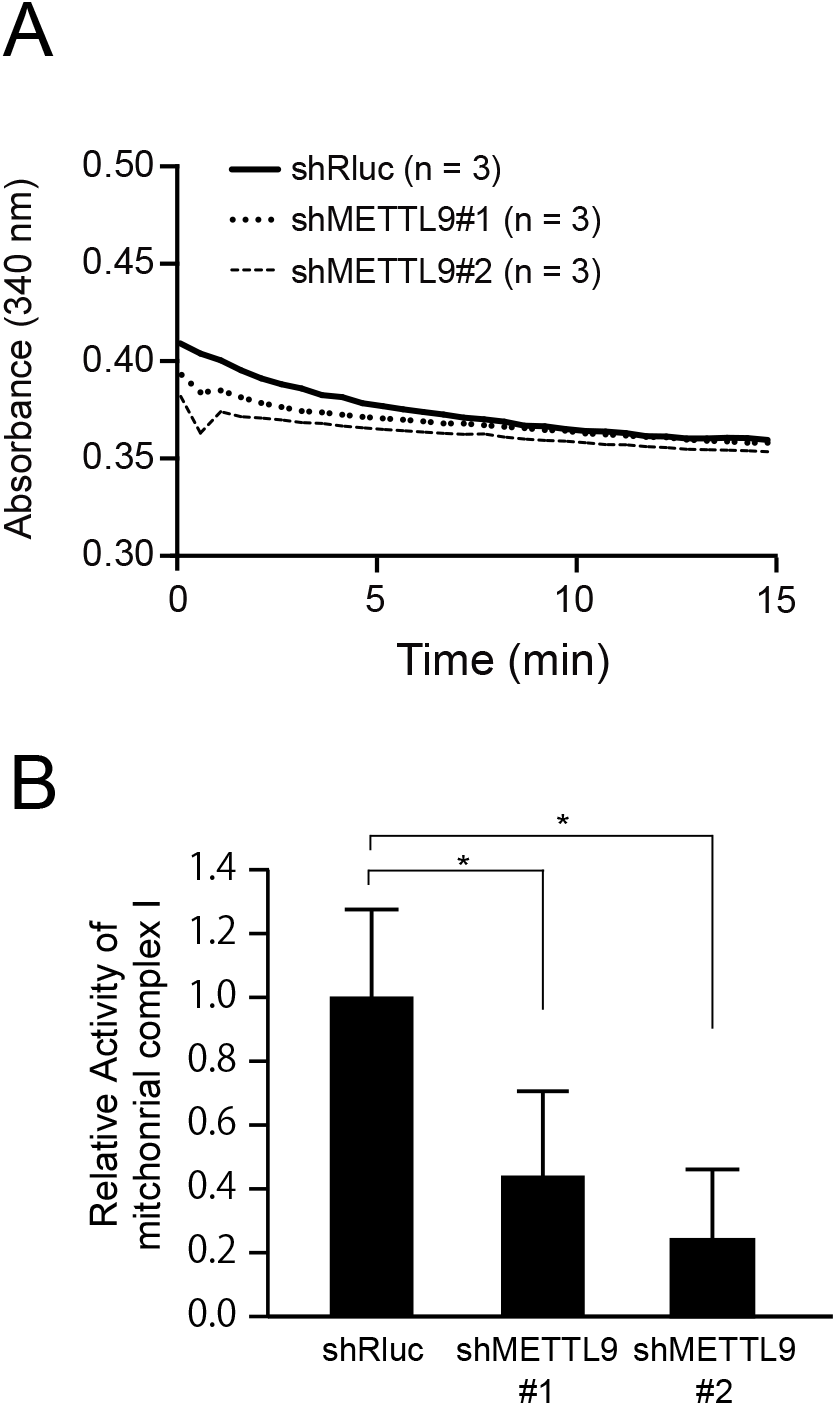
Knockdown of METTL9 in 58As9 cells reduces mitochondrial Complex I activity. (A) Mitochondrial Complex I activity was assessed using mitochondria isolated from METTL9 knockdown 58As9 cells. The knockdown showed the reduced activity of mitochondrial Complex I compared with control cells (shRluc). (B) The rate of mitochondrial Complex I activity was calculated by the slope of the linear portion of the curve. Knockdown of METTL9 showed significant reduction of mitochondrial Complex I activity. *, p<0.05.

## 4. Discussion

First, the present findings indicate that elevated METTL9 expression is closely associated with peritoneal dissemination in human scirrhous gastric cancers, as was clear by the evidence obtained from our metastatic cell experiments (Figs. 1, 2, and Supplementary Fig. 1). Further, using clinical tissue specimens of gastric cancers, we showed that METTL9 elevation is characteristic in patients with peritoneal dissemination, indicating that METTL9 can be a promising target molecule for inhibiting peritoneal dissemination. Because elevated METTL9 is observed in both primary cancer tissues and disseminated tissues, we are now vigorously examining METTL9 expression in circulating tumor cells in the bloodstream in gastric cancer patients with peritoneal dissemination. The present study is the first to connect elevated METTL9 with cancer metastasis.

Second, we found that METTL9 protein is predominantly localized in mitochondria in our metastatic scirrhous gastric cancer cells (Fig. 3). The localization pattern was similar to those in other reports showing the interaction of METTL9 with mitochondrial components [15, 16]. Further, a recent study found that METTL9 mediates pervasive 1-methylhistidine modification of NADH dehydrogenase (ubiquinone) 1 beta subcomplex 3 (NDUFB3) [15]. NDUFB3 is a subunit of mitochondrial respiratory Complex I, which activates mitochondrial respiration [15]. Our study also showed the reduced activity of mitochondrial respiratory Complex I in METTL9-knockdown cells (Fig. 4). Thus, our results suggest that METTL9-mediated 1-methylhistidine modification on the mitochondrial component is associated with the activity of mitochondrial Complex I. Dysregulation of mitochondria causes dynamic changes in intracellular energetic production and metabolic profiling, and thus supports the aggressiveness of cancer cells, i.e., cancer metastasis [17, 18]. Therefore, elevated METTL9 logically contributes to cancer cells’ acquisition of metastatic activity. In other words, inhibition of METTL9 activity can provide an effective therapeutic strategy against cancer metastasis.

The functions of the METTL family genes have attracted a great deal of attention. One member the family, METTL3, mediates the generation of N^6^-methyladenosine (m^6^A, nucleoside methylation). In contrast, METTL9 generates methylhistidine modified protein (protein methylation). Thus, each METTL protein catalyzes methylation for a unique target molecule. Such diversity of METTL functions makes it possible to regulate cell homeostasis, suggesting that abnormal expression of a METTL gene can be a trigger for cancers. A recent report showed that METTL9-mediated methylhistidine modification on zinc transporter SLC39A7 regulates the growth of prostate cancers both in vitro and in vivo [19]. When taken together, these reports indicate the interesting significance of the METTL family in cancers.

Metastasis-initiating cells (MICs) are a special subpopulation of cancer stem cells derived from primary tumors [20–24]. In concept, MICs are special cells that aggressively initiate cancer metastasis. According to the MIC concept, our metastatic cells (44As3 and 58As9) are regarded as peritoneal metastasis-initiating cells. They must have characteristics that are advantageous for cancer metastasis, and elevated METTL9 is one characteristic of MICs. Thus, METTL9 is a specific marker protein that arrests MICs in the bloodstream in gastric cancer patients.

## Conclusion

This study showed for the first time that METTL9 plays a significant role in peritoneal dissemination of human scirrhous gastric cancer. Our findings demonstrated that METTL9 is a suitable molecular target for the strategy of inhibiting peritoneal dissemination of scirrhous gastric cancer. METTL9 can be a specific marker to arrest peritoneal metastasis-initiating cells.

We hope that our findings can be applied to other types of cancers. This is a crucial starting point to study the significance of METTL family genes in cancer metastasis.

## Acknowledgements

We thank Dr. Ayami Morita-Kondo for her helpful suggestions regarding the experiments. We also thank Ms. Naomi Maruyama for her excellent technical assistance. This work was supported in part by a Grant-in-Aid for Challenging Exploratory Research (15K15063), by Grants-in-Aid for Scientific Research B (16H04697 and 19H03149), and by a Grant-in-Aid for Young Scientists B (17K14989) from the Japan Society for the Promotion of Science. We also express our gratitude for grant support from the Institute of Pharmaceutical Life Sciences, Aichi Gakuin University.

## Supplementary Materials

### Supplementary Figure Legend

**Supplementary Fig 1.**
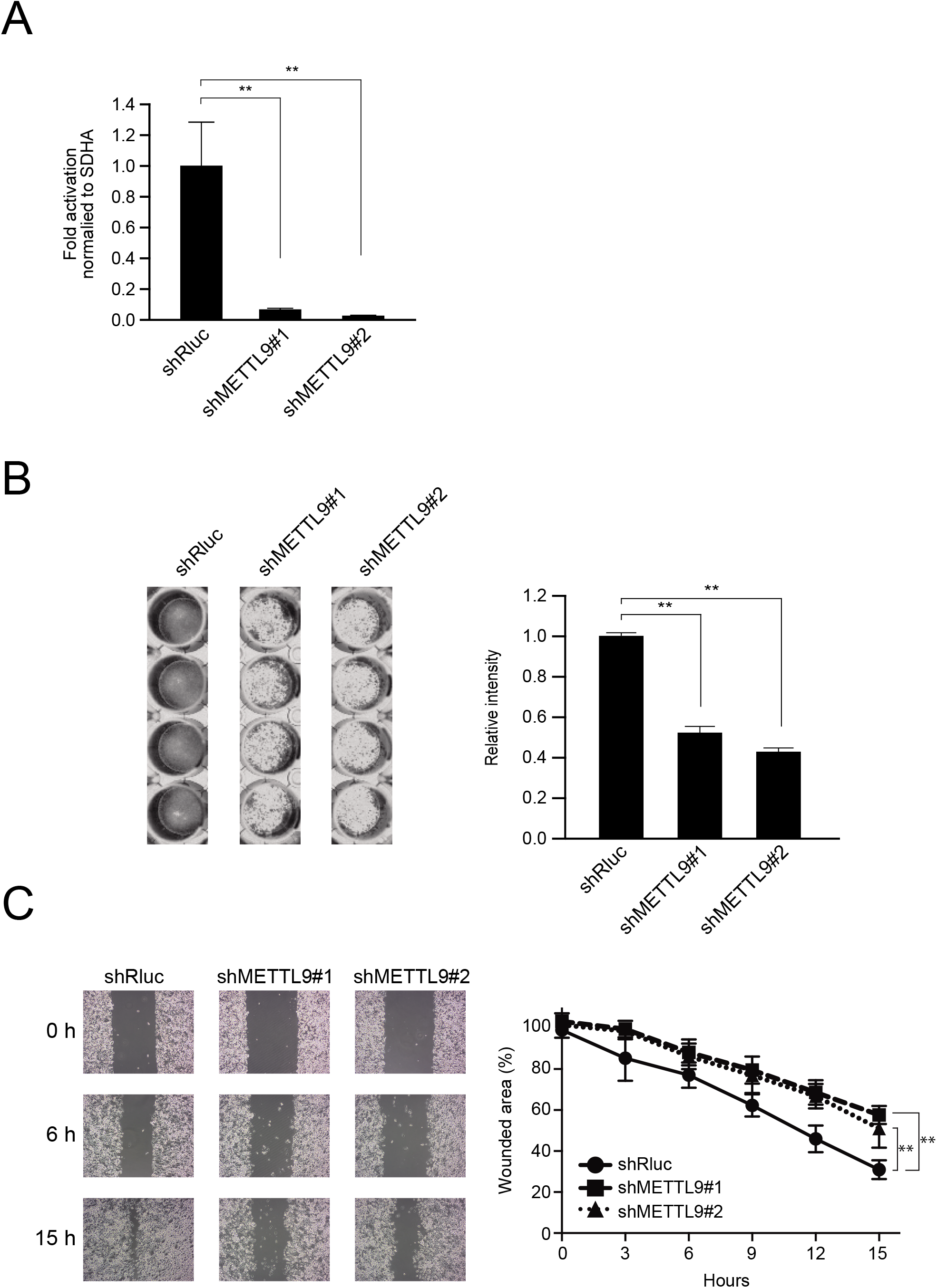
Similar phenotypic effects were obtained in METTL9-knockdown in 44As3 metastatic cells established from another scirrhous gastric cancer patient. (A) Knockdown efficiencies of shRNA targeting METTL9 in 44As3 were examined by RT-qPCR. Both shMETTL9#1 and shMETTL9#2 showed significant reductions in METTL9 expression compared with shRluc. **, p<0.01. (B) Clonogenic assay. After crystal violet staining, the images were obtained. The pixel intensity of each image was analyzed by Image J software, as shown in the right panel in (B). Significantly fewer colonies of METTL9 knockdown 44As3 cells were formed compared with the control cells (shRluc). **, p<0.01. (C) Cell migration assay (scratch assay). The effect of METTL9 knockdown 44As3 metastatic cells on cell migration was examined by the wound-healing assay method. Knockdown of METTL9 in 44As3 cells revealed significant inhibition of cell migration compared with the control cells (shRluc). **, p<0.01.

## Supplementary Methods

### Clonogenic assay

Five-hundred cells were seeded in a 24-well plate. After 10 days, the cells were fixed with ice-cold methanol for 10 minutes and then stained with 0.5% crystal violet solution containing 20% methanol for 30 min. The cells were washed with distilled water 3 times and placed at room temperature until completely dried. The picture was taken with an LAS 4000 digital imaging system (GE Healthcare).

### Wound-healing assay

Cells (5 × 10^5^) were seeded in a culture insert (iBidi) placed onto a 24-well plate. After 24 hours, the culture insert was removed and then a picture of the wounded areas was taken through a microscope every 3 hours. The wounded region was calculated by ImageJ software with the MRI Wound Healing Tool.

---

**Supplementary Table 1.**
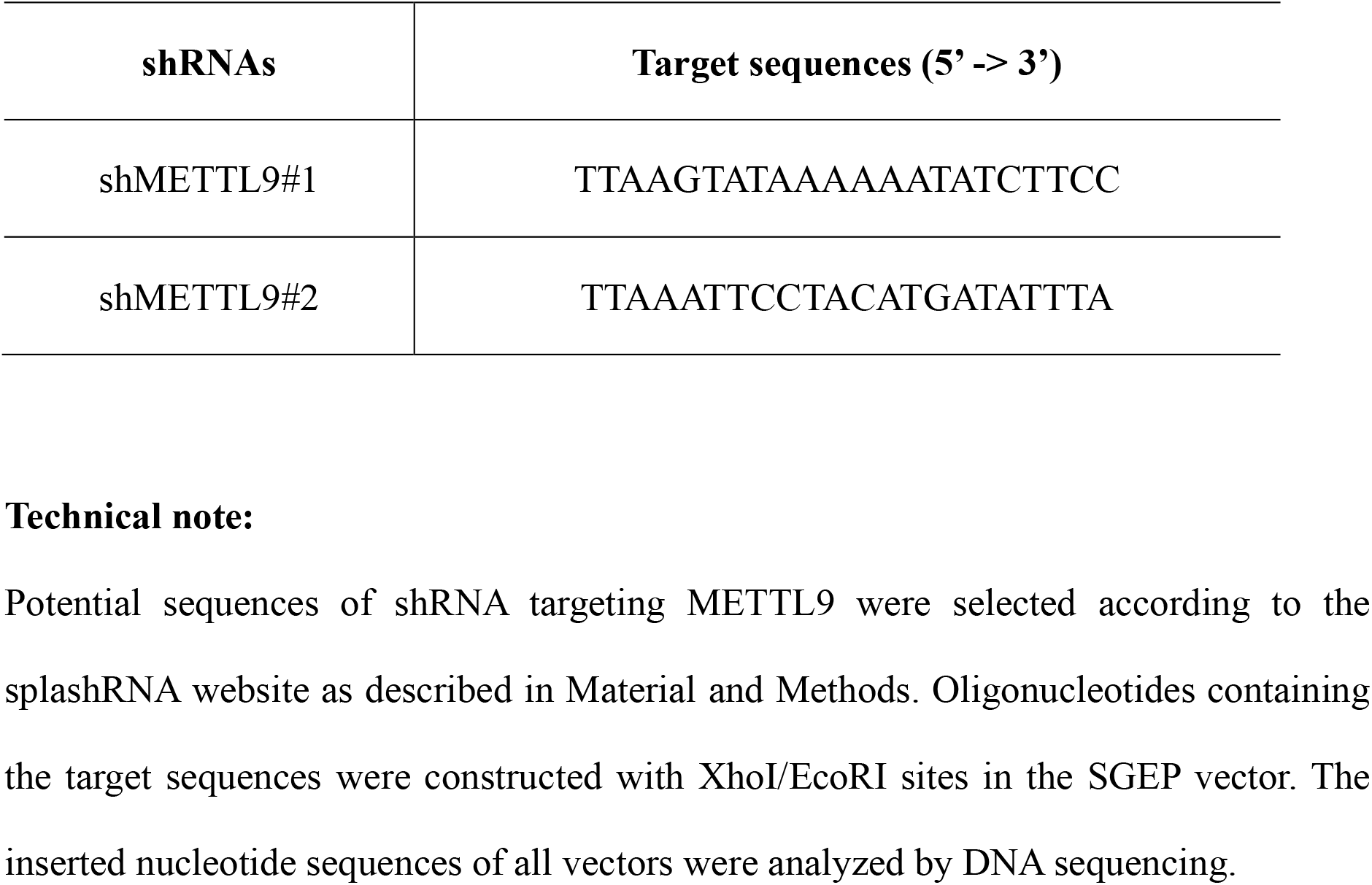
Target sequences of shRNAs for human METTL9.

**Supplementary Table 2.**
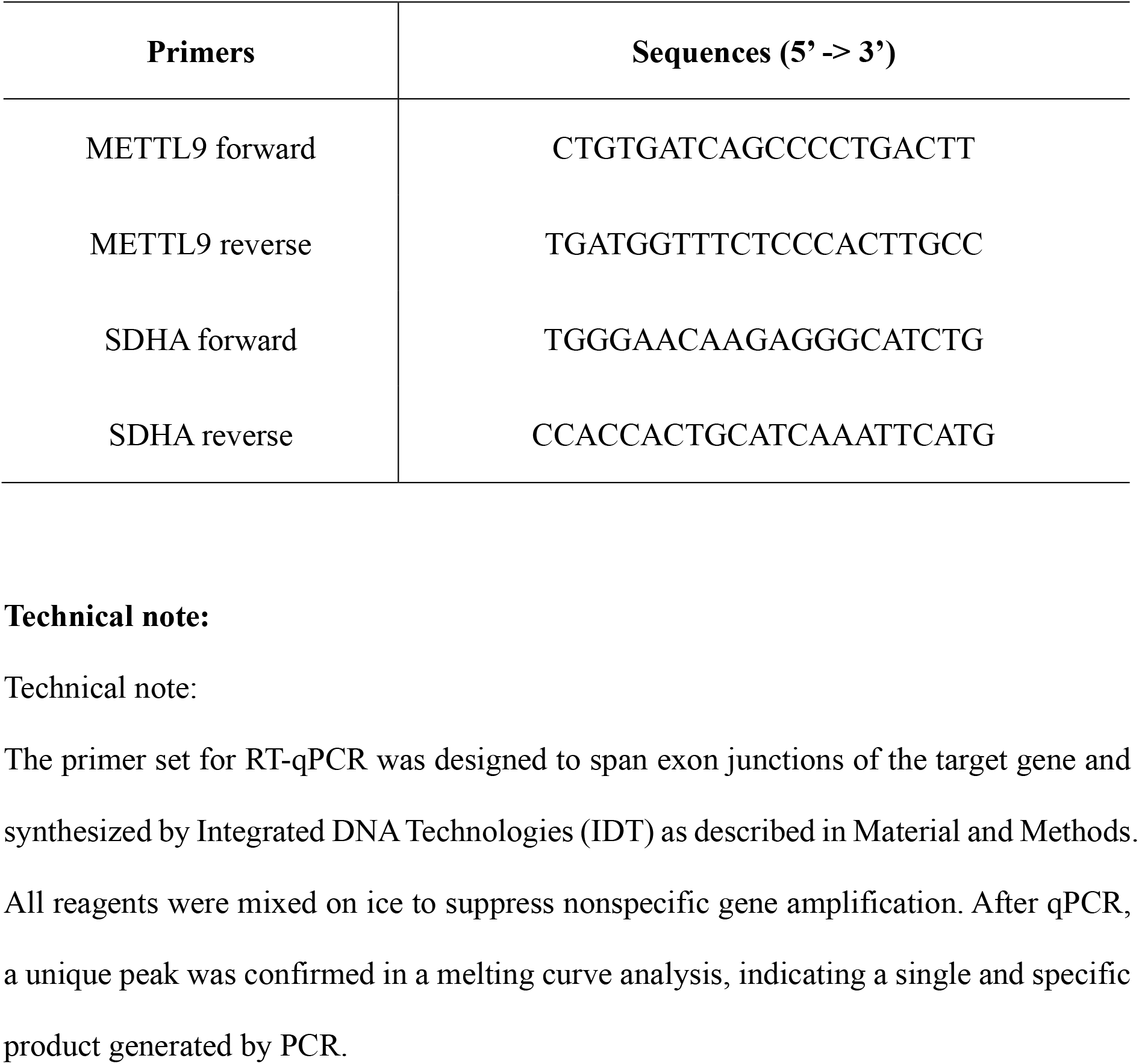
Specific primer sequences used for RT-qPCR.

